# Identification of a Proteomic Signature of Senescence in Primary Human Mammary Epithelial Cells

**DOI:** 10.1101/2020.09.22.309351

**Authors:** Alireza Delfarah, DongQing Zheng, Jesse Yang, Nicholas A. Graham

**Affiliations:** Mork Family Department of Chemical Engineering and Materials Science, University of Southern California, Los Angeles, CA 90089; Norris Comprehensive Cancer Center, University of Southern California, Los Angeles, CA 90089; Leonard Davis School of Gerontology, University of Southern California, Los Angeles, CA 90089

**Keywords:** replicative senescence, aging, mammary epithelial cell, proteomics, data integration, transcriptomics

## Abstract

Senescence is a permanent cell cycle arrest that occurs in response to cellular stress. Because senescent cells promote age-related disease, there has been considerable interest in defining the proteomic alterations in senescent cells. Because senescence differs greatly depending on cell type and senescence inducer, continued progress in the characterization of senescent cells is needed. Here, we analyzed primary human mammary epithelial cells (HMECs), a model system for aging, using mass spectrometry-based proteomics. By integrating data from replicative senescence, immortalization by telomerase reactivation, and drug-induced senescence, we identified a robust proteomic signature of HMEC senescence consisting of 77 upregulated and 36 downregulated proteins. This approach identified known biomarkers, such as downregulation of the nuclear lamina protein lamin-B1 (LMNB1), and novel upregulated proteins including the β-galactoside-binding protein galectin-7 (LGALS7). Gene ontology enrichment analysis demonstrated that senescent HMECs upregulated lysosomal proteins and downregulated RNA metabolic processes. We additionally integrated our proteomic signature of senescence with transcriptomic data from senescent HMECs to demonstrate that our proteomic signature can discriminate proliferating and senescent HMECs even at the transcriptional level. Taken together, our results demonstrate the power of proteomics to identify cell type-specific signatures of senescence and advance the understanding of senescence in primary HMECs.

## Introduction

Cellular senescence is a complex stress response that results in permanent cell cycle arrest. Multiple stressors can induce senescence, including replicative stress (e.g., telomere attrition), DNA damage (e.g., DNA double strand breaks), reactive oxygen species, oncogene activation, and drug-induced stress^1^. Senescence plays a role in development^2,3^ and wound healing^4,5^ but is most famous as a protective stress response against cancer^6^. However, as senescent cells accumulate in aging tissues^7^, they drive multiple age-related pathologies including atherosclerosis^8^, cardiac dysfunction^9^, diabetes^10^, kidney dysfunction^11^, osteoarthritis^12^, and overall decrements in healthy lifespan^13,14^. Many of the pro-disease effects of senescence are mediated by the senescence-associated secretory phenotype (SASP), a complex mixture of cytokines secreted by senescent cells that promote inflammation, wound healing, and growth responses in nearby cells^15,16^. In mice, therapies that specifically target senescent cells have shown great promise in the prevention or attenuation of age-related disease^13,14,17,18^.

Given their role in age-related disease, the characterization of senescent cells is important, particularly because biomarkers for senescence differ from cell type to cell type. The cell type-specificity of senescence biomarkers is perhaps not surprising given that the molecular mechanisms underlying senescence and SASP depend on the senescence inducer and cell type^19,20^. Despite considerable progress in our understanding of senescence, the molecular mechanisms underlying cellular senescence have not been fully elucidated^21–23^ and continued progress in the characterization of senescent cells is needed to improve our understanding of aging and disease.

Proteomics has emerged as a powerful tool for the characterization of senescent cells including plasma membrane-associated biomarkers^24^, proteomic alterations in the aging lung^25^, the therapy-induced senescence proteome^26^, and characterization of SASP^19^. In this study, we used liquid chromatography-mass spectrometry (LC-MS)-based proteomics to characterize the proteome of senescent primary human mammary epithelial cells (HMECs). These primary cells have been previously shown to accurately represent the molecular changes that occur during replicative senescence *in vivo*^27^. By integrating proteomic data across three data sets, we identified a core HMEC senescence signature of 77 upregulated and 36 downregulated proteins. This signature included both well-characterized senescence biomarkers (e.g., downregulation of lamin B1) and novel upregulation proteins in senescent cells (e.g., galectin-7). By integration of our proteomic signature of HMEC senescence with transcriptomic data from senescent HMECs, we further demonstrated that our proteomic signature is broadly recapitulated at the transcriptional level and can discriminate proliferating and senescent HMECs using transcriptional data. Taken together, our results demonstrate the power of proteomics to identify cell type-specific signatures of senescence and advance the understanding of senescence in primary HMECs.

## Experimental Procedures

### Cell culture

Primary HMEC cells were purchased from Thermo Scientific and cultured in M87A medium (50% MM4 medium and 50% MCDB170 supplemented with 5 ng/ml EGF, 300 ng/ml hydrocortisone, 7.5 μg/ml insulin, 35 μg/ml BPE, 2.5 μg/ml transferrin, 5 μM isoproterenol, 50 μM ethanolamine, 50 μM o-phosphoethanolamine, 0.25 % FBS, 5 nM triiodothyronine, 0.5 nM estradiol, 0.5 ng/ml cholera toxin, 0.1 nM oxytocin, 1% anti-anti, no AlbuMax) in atmospheric oxygen^28^. Glucose and glutamine-free DMEM was purchased from Corning (90-113-PB), Ham’s F12 was purchased from US Biological (N8542-12), and MCD170 medium was purchased from Caisson Labs (MBL04). Glucose and glutamine were added to the media at the appropriate concentration for each media type. Cells were lifted with TrypLE at 80-90% confluency and seeded at a density of 2.3 × 10^3^/cm^2^. To genetically modify HMECs, proliferating HMECs were infected at PD 14 with pLenti-PGK-hygro (Addgene 19066) encoding either hTERT or firefly luciferase. Following infection, cells were selected with 5 μg/ml hygromycin for 7 days. Following selection, cells were maintained in culture with 2 μg/ml hygromycin. To pharmacologically induce senescence, proliferating HMECs at PD ~8 were treated with 2 μM of the RRM2 inhibitor triapine or DMSO for 72 h. All samples profiled by LC-MS proteomics were collected in parallel with the characterization of senescence biomarkers as described in our previous publication^29^.

### LC-MS proteomics

Cell culture dishes were placed on ice and washed with PBS. Cells were then scraped and pelleted by centrifugation. The cell pellets were lysed by probe sonication in 8 M urea (pH 7.5), 50 mM Tris, 1 mM activated sodium vanadate, 2.5 mM sodium pyrophosphate, 1 mM β-glycerophosphate, and 100 mM sodium phosphate. The above procedures were performed in 0-4°C. Insoluble cell debris were filtered by 0.22 μm syringe filter. Protein concentration was measured by BCA assay (Pierce, PI23227). Lysates were reduced with 5 mM DTT, alkylated with 25 mM iodoacetamide, quenched with 10 mM DTT, and acidified to pH 2 with 5% trifluoracetic acid. Proteins were then digested to peptides using a 1:100 trypsin to lysate ratio by weight. Tryptic peptides were desalted by reverse phase C18 StageTips and eluted with 30% acetonitrile. The eluents were vacuum dried, and 250 ng/injection was submitted to LC-MS. Samples were randomized and injected into an Easy 1200 nanoLC ultra high-performance liquid chromatography coupled with a Q Exactive Plus quadruple orbitrap mass spectrometry (Thermo Fisher). Peptides were separated by a reverse-phase analytical column (PepMap RSLC C18, 2 μm, 100Å, 75 μm × 25 cm). Flow rate was set to 300 nL/min at a gradient from 3% buffer B (0.1% formic acid, 80% acetonitrile) to 38% B in 110 min, followed by a 10-min washing step to 85% B. The maximum pressure was set to 1,180 bar and column temperature was maintained at 50°C. Peptides separated by the column were ionized at 2.4 kV in the positive ion mode. MS1 survey scans were acquired at the resolution of 70k from 350 to 1800 m/z, with maximum injection time of 100 ms and AGC target of 1e6. MS/MS fragmentation of the 14 most abundant ions were analyzed at a resolution of 17.5k, AGC target 5e4, maximum injection time 65 ms, and normalized collision energy 26. Dynamic exclusion was set to 30 s and ions with charge +1, +7, and >+7 were excluded. MS/MS fragmentation spectra were searched with Proteome Discoverer SEQUEST (version 2.2, Thermo Scientific) against in-silico tryptic digested Uniprot all-reviewed *Homo sapiens* database (release Jun 2017, 42,140 entries) plus all recombinant protein sequences used in this study. The maximum missed cleavages was set to 2. Dynamic modifications were set to oxidation on methionine (M, +15.995 Da) and acetylation on protein N-terminus (+42.011 Da). Carbamidomethylation on cysteine residues (C, +57.021 Da) was set as a fixed modification. The maximum parental mass error was set to 10 ppm, and the MS/MS mass tolerance was set to 0.02 Da. The false discovery threshold was set to 0.01 using the Percolator Node validated by q-value. The relative abundance of parental peptides was calculated by integration of the area under the curve of the MS1 peaks using the Minora LFQ node. The RAW and processed LC-MS files have been uploaded to the PRIDE database ^30^ (PXD019057, Username, Reviewer password: djc8bohx).

### Data processing and normalization

Missing peptide abundances were imputed using the K-nearest neighbor algorithm^31^. The optimized number of neighbors was determined to be n = 10. Protein abundance log_2_ ratios and statistical significance were calculated using DEqMS in R software^32^. Briefly, peptide sequences were aggregated into protein log_2_ ratios by the median sweeping method: raw intensity values were log_2_ transformed, the median of log_2_ intensity was subtracted for each PSM, and then for each protein, the relative log_2_ ratio was calculated as the median of log_2_ ratio of the PSMs assigned to that protein. To calculate the statistical significance across the three datasets, individual p values were combined using Fisher’s Method and then corrected for multiple hypothesis testing using the Benjamini-Hochberg method.

### Hierarchical clustering

Clustering was performed using Morpheus from the Broad Institute (https://software.broadinstitute.org/morpheus/). Data was centered for each individual experiment prior to clustering with the metric one minus the Pearson correlation coefficient.

### Gene Ontology Enrichment Analysis

Gene ontology (GO) enrichment analysis was conducted in ClueGO^33^. Proteins with FDR-corrected p value less than 0.05 and either upregulated or downregulated log_2_ fold change greater than log_2_(1.5). Pathways included were GO biological process, GO cellular component, GO molecular function, and KEGG. The list of 907 quantified proteins was used as a custom background set, and p values were calculated with a right-tailed hypergeometric test and corrected for multiple hypothesis testing using the Bonferroni step down method.

### Transcription Factor Targets Enrichment Analysis

Proteins were ranked by their log_2_ (senescent / proliferating) fold change values. Gene set enrichment analysis (GSEA)^34^ was run with the unweighted statistic using the GSEA java applet using Broad Institute C3 TFT:GTRD (Gene Transcription Regulation Database) gene sets^35^.

### HMEC Senescence score

Log_2_-transformed, RMA-normalized Entrez gene expression values from pre-stasis, intermediate, and stasis HMECs^36^ were obtained for each individual cell line and media type. The HMEC senescence signature used for voting weights was created using proteins with absolute average log_2_ (senescent / proliferating) fold change greater than 1 and FDR-corrected p-value less than 0.01 from the combined HMEC proteomic analysis (Fig. 2). HMEC senescence scores were calculated by multiplying the senescence signature matrix versus the gene expression matrix. For the receiver operating characteristic (ROC) curve, HMEC senescence scores were used from pre-stasis and stasis samples only. The area-under-the-curve (AUC) was calculated by the composite trapezoidal rule.

## Results and Discussion

### Replicative senescence alters the HMEC proteome

To identify proteomic changes that accompany replicative senescence, we analyzed primary human mammary epithelial cells (HMECs) using quantitative, label-free LC-MS-based proteomics. We have previously found that primary HMECs enter senescence at ~35 population doublings (PD) and exhibit molecular markers of senescence including upregulation of senescence-associated β-galactosidase (SA-β-gal), upregulation of the cell cycle inhibitor p21, and cessation of DNA synthesis^29^. For this study, samples were collected at PD 8/9 (proliferating) or PD 35/36 (senescent) in parallel with the characterization of senescence biomarkers published in our previous study (Supp. Fig. 1A). Comparing proliferating and senescent HMECs with LC-MS proteomics, we quantified 1,303 proteins in two independent biological replicates (Supp. Fig. 1B and Supp. Table 1). Of these proteins, 60 were significantly upregulated and 39 were significantly downregulated in senescent HMECs (FDR-corrected p-value < 0.01 and average absolute log_2_ fold change > 1) (Fig. 1A). Hierarchical clustering of the individual sample values for significantly changing proteins demonstrated high reproducibility across biological and technical replicates (Fig. 1B). Among the most upregulated proteins in senescent cells was annexin A1 (ANXA1), which is associated with aging in the rat prostate^37^. Additionally, the β-galactosidase GLB1, which is associated with the senescence biomarker SA-β-gal activity^38^, was significantly upregulated in senescent HMECs. Significantly downregulated proteins in senescent HMECs included histone H4 (HIST1H4A) and SLC3A2 (also known as 4F2), a component of several heterodimeric amino acid transporter complexes including the cystine-glutamate antiporter xCT. Taken together, our proteomic profiling demonstrated that replicative senescence significantly changed the HMEC proteome.

**Figure 1:**
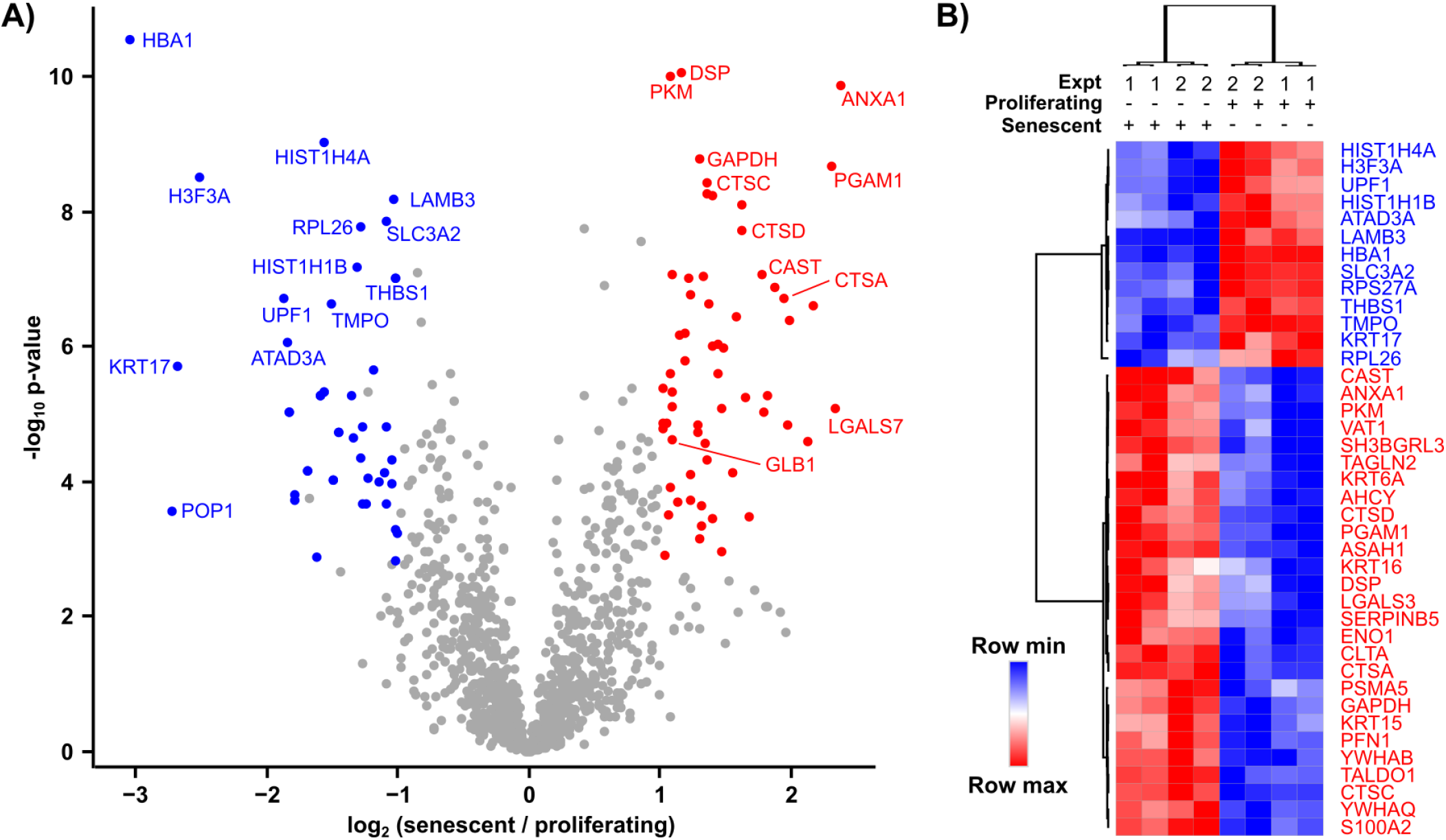
Replicative senescence alters the HMEC proteome. A. Volcano plot representing average log_2_ fold change of protein levels comparing replicative senescent versus proliferating HMECs plotted against the -log10 p-value (n=2 biological replicates). Red and blue denote the 60 significantly upregulated and 39 significantly down-regulated proteins, respectively (FDR-corrected p-value < 0.01 and average absolute log_2_ fold change > 1). B. Hierarchical clustering of protein expression levels for differentially expressed proteins in senescent and proliferating HMECs across two biological replicates (Expt.). For each biological replicate, the samples were run in technical replicates. The 40 proteins with smallest FDR-corrected p-values are shown.

**Figure 2:**
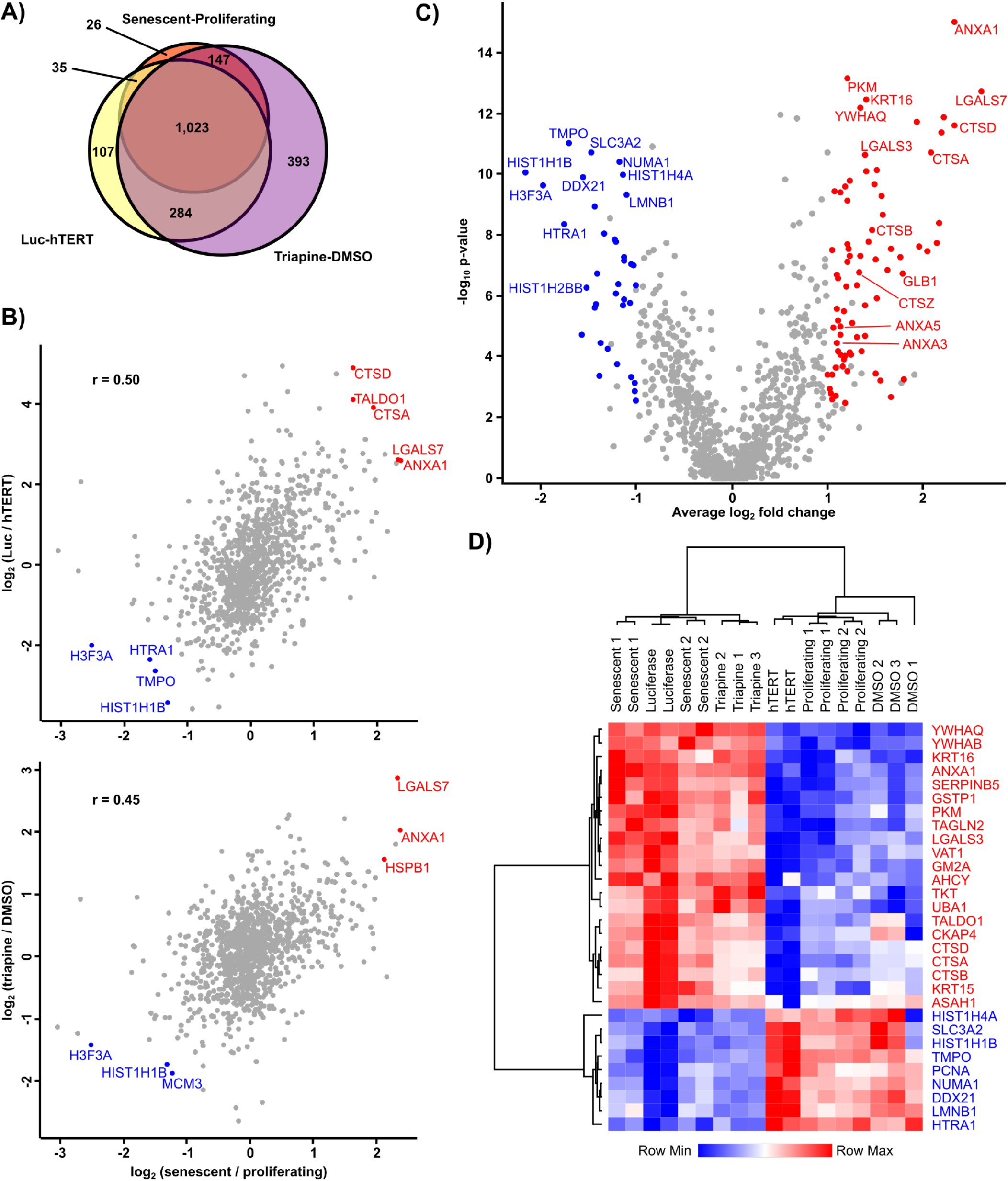
Data integration identifies a proteomic signature of HMEC senescence. A. Venn diagram showing the overlap in the number of proteins quantified in three senescent datasets: replicative senescent versus proliferating HMECs (Fig. 1), luciferase-versus hTERT-expressing HMECs (Supp. Fig. 2A,B), and triapine-versus DMSO-treated HMECs (Supp. Fig. 2C,D). B. Comparison of log_2_ fold change in protein expression in luciferase-versus hTERT-expressing HMECs (top) and triapine-versus DMSO-treated HMECs (bottom) against replicative senescent versus proliferating HMECs. The Pearson correlation coefficient (r) is shown. C. Volcano plot representing average log_2_ (senescent / proliferating) fold change versus -log10 p-value combined statistical significance from three datasets. Red and blue denote 76 and 36 proteins that were consistently up- or down-regulated in all three datasets, respectively (FDR-corrected p-value < 0.01, average absolute log_2_ fold change > 1). D. Hierarchical clustering of the 30 proteins with the smallest FDR-corrected p-value after integrating the three datasets. All biological and technical replicates are shown.

### Data integration identifies a proteomic signature of HMEC senescence

Next, we sought to identify a core signature of HMEC senescence by integrating proteomic data from replicative senescence (Fig. 1) with proteomic profiling of HMECs that had been manipulated to either bypass senescence or to prematurely senesce. We leveraged our previous findings that 1) expression of human telomerase reverse transcriptase (hTERT) immortalizes HMECs and enables bypass of replicative senescence and that 2) inhibition of the nucleotide synthesis enzyme RRM2 with the drug triapine induces premature senescence in proliferating HMECs^29^. We then used label-free LC-MS proteomics to compare 1) hTERT-immortalized HMECs (i.e., proliferating) to senescent HMECs expressing the negative control protein luciferase and 2) proliferating HMECs treated for 3 days with either DMSO (i.e., proliferating) or triapine (i.e., senescent) (Supp. Fig. 2 and Supp. Tables 2 and 3). Again, these samples were collected in parallel with our previously published characterization of senescence biomarkers^29^. Across all three datasets, LC-MS proteomics consistently quantified 1,023 proteins in all biological and technical replicates (Fig. 2A and Supp. Table 4). Comparing the hTERT/luciferase and DMSO/triapine proteomic signatures to that of replicative senescence, we found that both signatures were significantly correlated with the proteomic changes observed in replicative senescence (Fig. 2B). Plotting the combined data on a volcano plot revealed 76 and 36 significantly upregulated and downregulated proteins, respectively (combined FDR-corrected p-value < 0.01 and average absolute log_2_ fold change > 1) (Fig. 2C). Hierarchical clustering of the individual biological and technical replicates demonstrated consistent upregulation or downregulation for the most significantly changing proteins across the three individual proteomic signatures (Fig. 2D). Thus, integration across the three data sets identified a robust proteomic signature of HMEC senescence.

Several proteins identified in the HMEC senescence proteomic signature are previously known senescence biomarkers. Among the significantly downregulated proteins was the nuclear lamina component lamin-B1 (LMNB1). Loss of lamin-B1 expression in senescent cells has been extensively documented, including in replicative senescence, oncogene-induced senescence, and UV-induced senescence^39–43^. Notably, the lamin-B1-binding partner TMPO (LAP2) was also part of our senescence signature, although decreases in TMPO expression are not unique to senescent cells, as downregulation also occurs in quiescent cells^40^. Regardless, the concordance of LMNB1 expression in our HMEC system and other studies adds additional support that loss of LMNB1 expression is a bona fide senescence biomarker. The most downregulated protein in our proteomic signature of HMEC senescence was the histone H1.5 (HIST1H1B) with an average log_2_ fold change in senescent cells of −2.16 (Fig. 2C). We additionally observed downregulation of five additional histone proteins in our combined proteomics analysis including H1.3 (HIST1H1D), H2A.Z (H2AFZ), H2B type 1-J (HIST1H2BJ), H2B type 2-F (HIST2H2BF), and H4 (HIST1H4A). Consistent with our findings, several studies have reported loss of histone H1 and DNA methylation in senescence and aging^44–46^. These results support the regulatory role of chromatin remodeling in HMEC senescence.

The most significantly upregulated protein in our HMEC senescence signature was the calcium-dependent phospholipid-binding protein annexin 1 (ANXA1) with an average log_2_ fold change in senescent cells of 2.33 (Fig. 4B). We also observed significant upregulation of two other annexins, ANXA3 and ANXA5, in senescent HMEC (average log_2_ fold change 1.1 for both proteins). Interestingly, the upregulation of annexins has been previously linked to increased lipid metabolism in a model of therapy-induced senescence^26^. Moreover, accumulation of nuclear ANXA5 is a biomarker of replicative and therapy-induced fibroblast senescence^47^, and secretion of ANXA1, ANXA3, and ANXA5 is upregulated in senescent fibroblasts^19^. In addition, we observed upregulation of several lysosomal proteins in senescent HMEC including the GLB1 (β-galactosidase), four cathepsins (CTSA, CTSD, CTSD, and CTSZ), and the glycosylase MAN2B1. These results are consistent with previous reports of increased lysosomal activity in senescence^48,49^. Additionally, cathepsins are known to regulate senescence^50^ and pathogenesis of age-related disease^48^ and are also secreted by senescent cells^19^. Finally, both the β-galactoside-binding proteins galectin-3 (LGALS3) and galectin-7 (LGALS7) were significantly upregulated in senescent HMECs. Galectin-3 can coordinate repair, removal, and replacement of lysosomes^51^, and its upregulation may reflect attempts by senescent cells to repair deteriorating lysosomes^52^. To our knowledge, galectin-7 has not been reported to be involved in senescence, but we speculate that it may also play a role in lysosomal repair and homeostasis in senescent HMEC. Taken together, these results suggest that annexins, cathepsins, and galectins are potential senescence biomarkers across many cell types.

### Gene ontology and transcription factor target analysis of the HMEC senescence signature

Next, to understand the functional classes of proteins altered upon replicative senescence, we performed gene ontology (GO) enrichment analysis. The most significantly upregulated GO terms in senescent HMECs included lysosome, antioxidant activity, and carbohydrate catabolic process (Fig. 3A). This result is consistent with the known upregulation of secretory pathways and lysosomes in senescence^53^. In addition, increased lysosomal activity has been linked to proteolysis of histones in senescent cells^54^, consistent with our observation that expression of several histones is decreased in senescent HMEC. The most significantly downregulated GO terms in senescent cells were ribosomal, translational, and RNA metabolism-related terms, consistent with reports that reduced RNA turnover and alterations in translation drive cellular senescence^55,56^. Together, this analysis suggests that our proteomic signature of HMEC senescence is broadly enriched for similar categories of proteins as other models of senescence.

**Figure 3:**
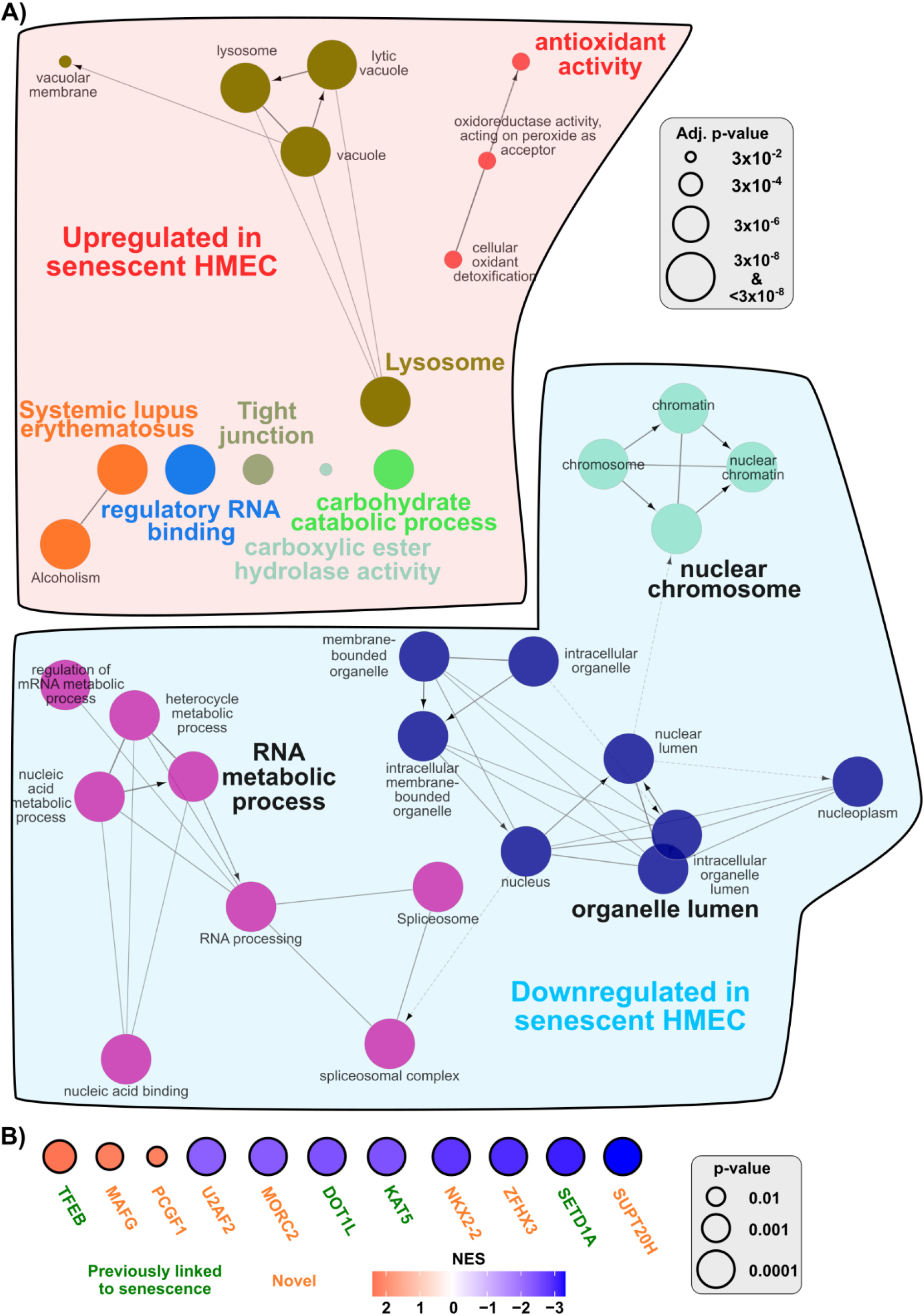
Gene ontology and transcription factor target analysis of the HMEC senescence signature. A. Gene Ontology (GO) enrichment analysis of the upregulated (top) and downregulated (bottom) proteins in the HMEC senescence signature. Analysis was performed in ClueGO^33^ using proteins with FDR-corrected p-value less than 0.05 and log_2_ fold change greater than log_2_(1.5). The size of each circle denotes the statistical significance of the enrichment, as shown in the legend. GO terms of the same color have ≥50% similarity. Connecting lines represent kappa connectivity scores. B. Transcription factor target analysis of proteomic signature of HMEC senescence. The 1,023 proteins quantified in all three proteomic data sets (Fig. 2) were ranked by average log_2_ fold change and analyzed by gene set enrichment analysis (GSEA) with the GTRD list of transcription factor targets^35^. Circle color denotes the normalized enrichment score (NES), and circle size denotes the statistical significance of the enrichment, as shown in the legend. Transcription factors with a known link to senescence are colored green, and transcription factors without any known link to senescence are colored orange.

**Figure 4:**
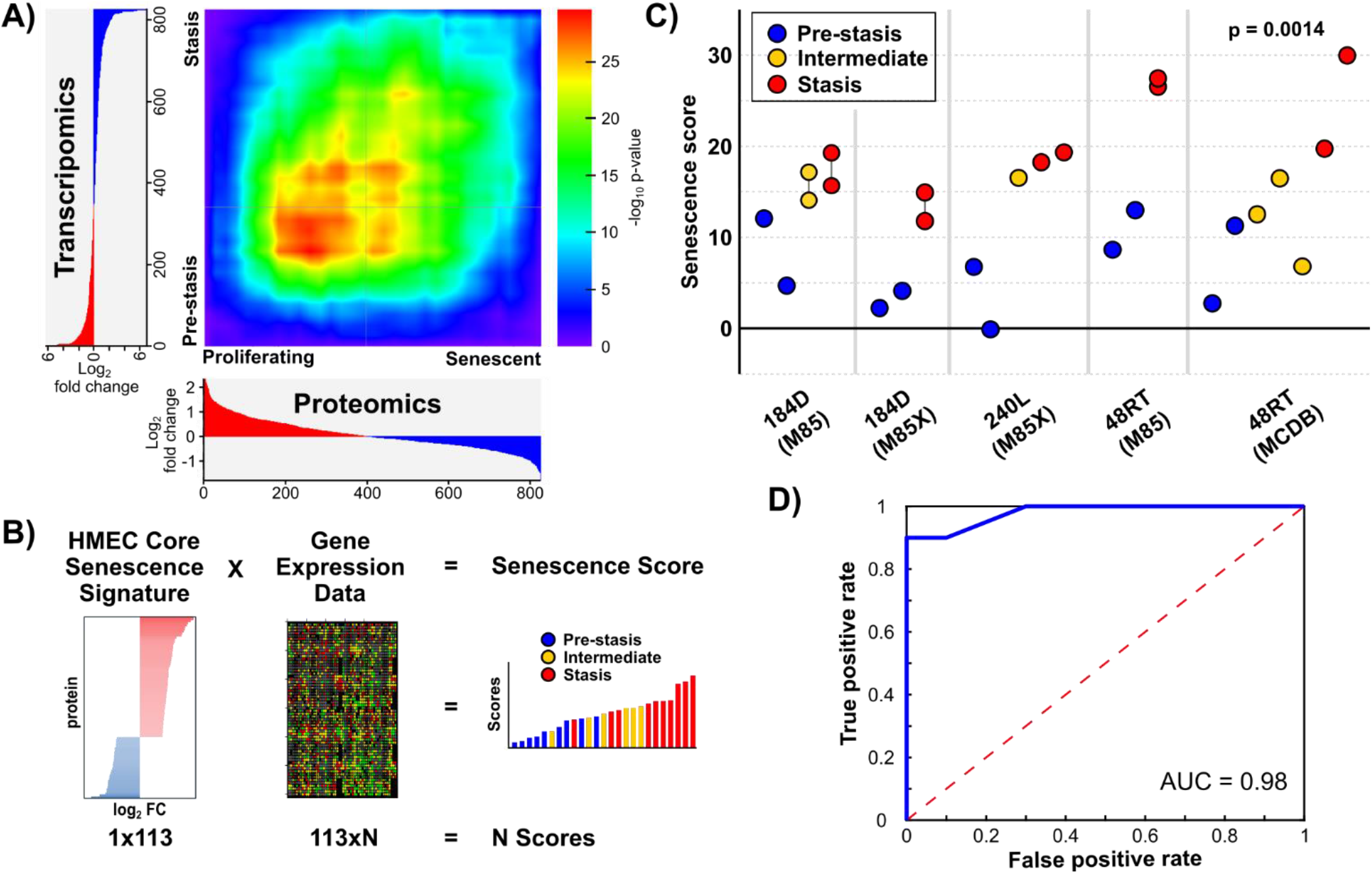
Classification power of the proteomic signature of HMEC senescence. A. Rank-rank hypergeometric overlap (RRHO)^61^ comparison of proteomic (Fig. 2) and transcriptomic signatures of HMEC senescence^36^. The protein and transcripts measured in both data sets (n=826) were ranked by log_2_ fold change. The area of most significant overlap, as shown by the color legend at right, occurs in the lower left-hand quadrant which are the proteins and transcripts upregulated in both data sets. B. Schematic representing calculation of HMEC senescence score using weighted voting^62^. The proteomic signature of HMEC senescence (Fig. 2) was used as voting weights (log_2_ fold change of protein expression comparing senescent and proliferating HMEC, 113 proteins total). Weights were multiplied by gene expression data to calculate a HMEC senescence score for each sample. C. Gene expression data from five independent HMEC cell lines^36^ was analyzed using weighted voting as described in panel B. Samples for each cell line are arranged in increasing passage number and colored according to pre-stasis (i.e., proliferating), intermediate, or stasis (i.e., senescent) as in the original publication. M85, M85X, and MCDB represent different media formulations. Samples profiled at the same passage are connected by a thin dark gray line. P = 0.0014 comparing the senescence scores from pre-stasis and stasis using a paired (by cell line) t-test. D. Receiver operating characteristic (ROC) curve analysis of the HMEC senescence score data in panel C. Only pre-stasis and stasis samples were included. The area-under-the-curve (AUC) was 0.98.

Next, to identify transcription factors that might regulate senescence, we performed enrichment analysis on the combined proteomics data using transcription factor target (TFT) gene lists^35^. This analysis identified 3 TFTs whose targets were significantly upregulated in senescent HMECs (TFEB, MAFG, PCGF1), and 40 TFTs whose targets were significantly downregulated in senescent HMECs including SUPT20H, SETD1A, and ZFKX3 (Fig. 3B, p-value < 0.05 and FDR q-value < 0.1). Several of the identified transcription factors have been previously linked to senescence and aging including downregulation of SETD1A^57^, KAT5^58^, and DOT1L^59^ as well as upregulation of TFEB^60^. In contrast, the transcription factors MAFG and PCG1 (targets upregulated) and NKX2-2, ZFHX3 and SUPT20H (targets downregulated) have not been previously linked to aging or senescence to our knowledge. Future studies are necessary to investigate whether these transcription factors are regulators of cellular senescence in HMECs and other cell types. Taken together, our TFT analysis identified both known and novel transcription factor regulators of senescence.

### Defining a senescence score that predicts HMEC senescence

Having identified an HMEC proteomic signature of senescence, we next asked whether proteomic changes were reflected at the transcriptional level in senescent HMECs. To answer this question, we turned to transcriptomic profiling data from pre-stasis HMECs (i.e., proliferating), intermediate HMECs, or HMECs at stasis (i.e., a stress-associated senescence barrier associated with elevated levels of p16 and/or p21, G1 arrest, and the absence of genomic instability)^36^. Analysis by rank-rank hypergeometric overlap (RRHO)^61^ showed significant overlap between our proteomic signature and that of HMEC stasis, particularly in the upregulated transcripts/proteins (Fig. 4A). We next asked whether our proteomic signature could discriminate pre-stasis and stasis samples in this transcriptional data set. We thus defined a weighted voting scheme^62^ where the log_2_ fold changes of the 113 core senescence proteins (Fig. 2) were multiplied by gene expression data from the same 113 genes. The result is a “senescence score” for each individual sample where increasing scores predict senescence (Fig. 4B). Testing this approach, we found that the senescence score was significantly increased for five independent HMEC cell lines as they entered stasis (Fig. 4C). The average increase in senescence score from pre-stasis to stasis was 3.6 ± 1.4 (p = 0.0014). Lastly, we plotted the HMEC senescence scores on a receiver operating characteristic (ROC) curve and found that the area-under-the-curve was 0.98 indicating high classification sensitivity and specificity (Fig. 4D). Together, these results indicate that our proteomic HMEC signature is broadly conserved at the transcriptional level in HMECs and can discriminate proliferating from senescent HMECs even at the transcriptional level.

## Conclusion

Cellular senescence is a state of irreversible cell cycle arrest that contributes to degenerative and hyperplastic phenotypes in aging, cancer, and many other diseases. Taken together, our combined analysis identified a core proteomic signature of HMEC senescence including potential novel senescence biomarkers and regulators. These results support that quantitative proteomics is a powerful approach to delineate cell-type specific signatures of senescence. Our results thus advance the understanding of senescence in primary HMECs and contribute to our understanding of how senescence impacts aging and disease.

## Acknowledgments

This work was supported by the Rose Hills Foundation, the 2020 AACR-Bayer Innovation and Discovery Grant (Grant Number 20-80-44-GRAH), the University of Southern California (USC) Provost’s Office, and the Viterbi School of Engineering.

## Author Contributions

AD and NAG designed research. AD, DZ, and JY performed research. AD, DZ, and NAG analyzed data. AD and NAG wrote the manuscript.

## Data Availability Statement

The RAW and processed LC-MS files have been uploaded to the PRIDE database^30^ (PXD019057, Username, Reviewer password: djc8bohx).

