## Supplemental Figures for "Identification of a Proteomic Signature of Senescence in Primary Human Mammary Epithelial Cells"

Running title: *Proteomic profiling of primary HMEC senescence*

To whom correspondence should be addressed: Nicholas A. Graham, University of Southern California, Los Angeles, 3710 McClintock Ave., RTH 509, Los Angeles, CA 90089. Phone: 213-240-0449;

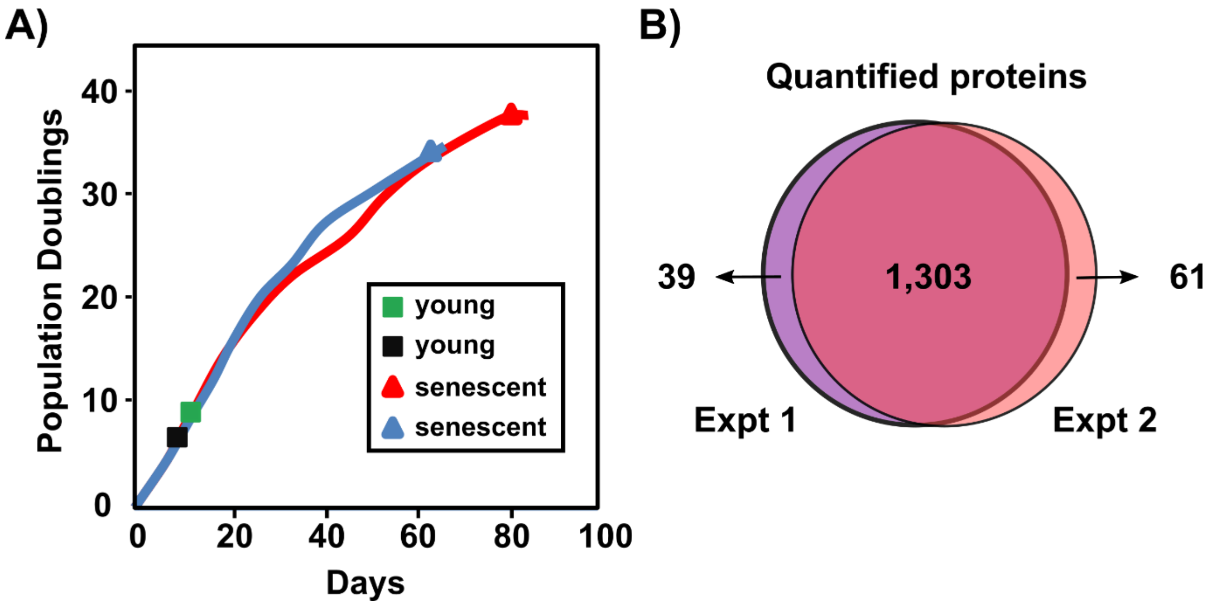


**Supporting Fig. S1: Sample collection and overlap of identified proteins for replicative senescence.**

A. Primary HMECs proliferate for ~35 population doublings (PD) before senescence due to replicative exhaustion. Proteomics samples were collected at PD 8/9 (proliferating) or PD 35/36 (senescent) in parallel with the characterization of senescence biomarkers published in our previous study[1], as shown on the growth curve.

B. Number of proteins quantified in two independent biological replicates as described in A. Proteomic analysis was performed as two independent biological replicates with one proliferating and one senescent sample in each experiment. For each experiment, duplicate injections (technical replicates) were run from each proliferating and senescent sample.

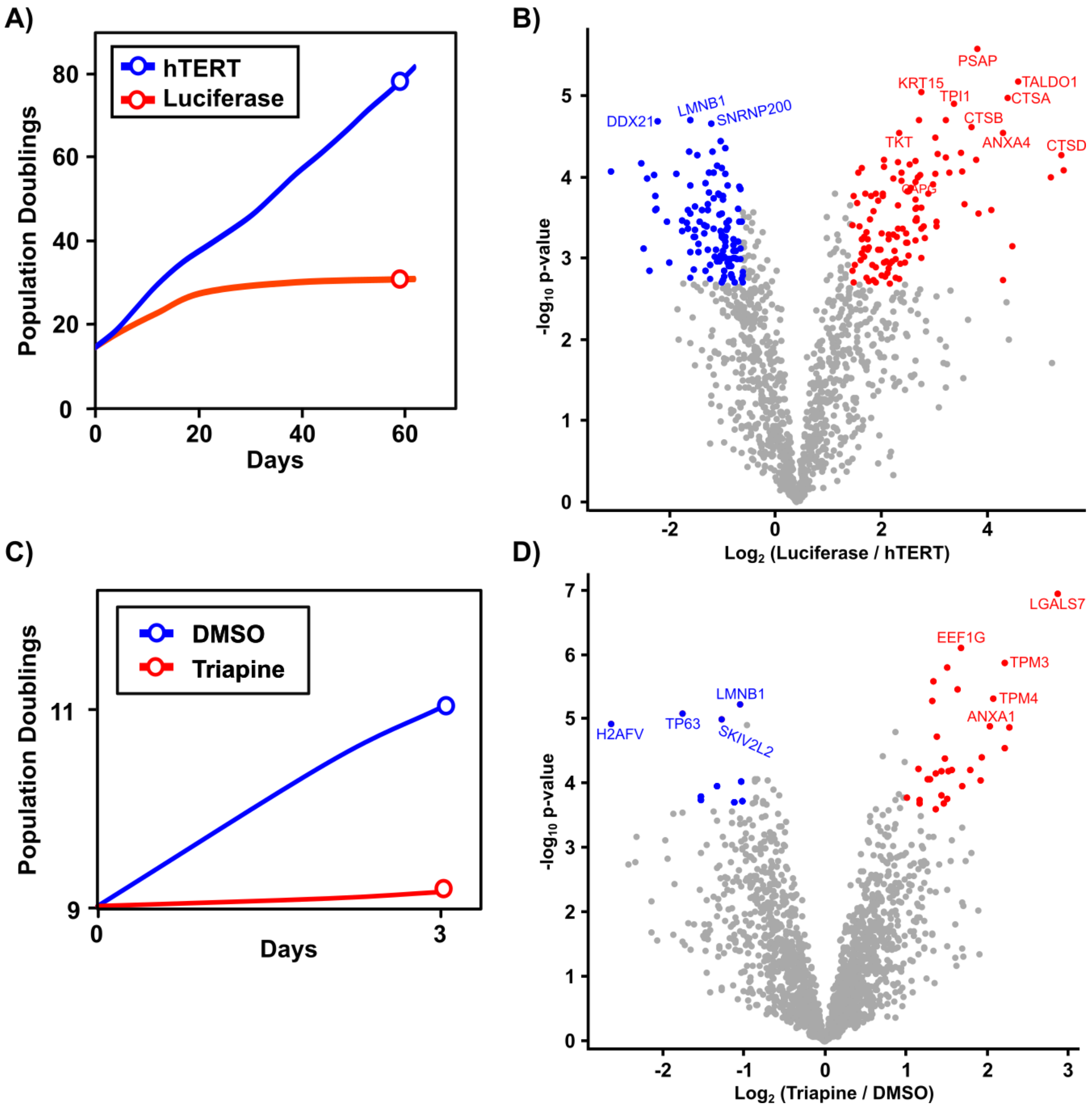


**Supporting Fig. S2: Analysis of hTERT-immortalized HMECs and triapine-induced senescence in HMECs**

A. Growth curve for HMECs infected with either luciferase (negative control, senescent) or hTERT (immortalized, proliferating). At ~60 days in culture, corresponding to 35 and 85 PD for luciferase and hTERT, respectively, hTERT-expressing HMECs continue to proliferate but luciferase-expressing HMECs are senescent. Circles represent the time of sample collection which was done in parallel with characterization of senescence biomarkers as described in our previous publication[1]. One biological replicate was collected and analyzed in technical duplicate by LC-MS proteomics.

B. Volcano plot representing $\log_2$ fold change of protein levels for luciferase versus hTERT plotted against the $-\log_{10}$ p-value. Red and blue denote significantly up- and down-regulated proteins, respectively (FDR-corrected p-value < 0.01 and absolute $\log_2$ fold change > 1). In total, we quantified 1,449 proteins (Supp. Table 2), of which 132 and 128 were significantly upregulated and downregulated, respectively, in senescent luciferase-expressing HMECs.

C. Growth curve for HMECs treated with either DMSO (negative control, proliferating) or triapine (senescent). Circles represent the time of sample collection. Characterization of senescence biomarkers was performed on the same cell lines as described in our previous publication[1]. Three biological replicates were collected and analyzed in technical singlicate by LC-MS proteomics.

D. Volcano plot representing $\log_2$ fold change of protein levels for triapine versus DMSO plotted against the $-\log_{10}$ p-value. Red and blue denote significantly up- and down-regulated proteins, respectively (FDR-corrected p-value < 0.01 and absolute $\log_2$ fold change > 1). In total, we identified 1,847 proteins (Supp. Table 3), of which 31 and 10 were significantly upregulated and downregulated, respectively, in triapine-treated senescent HMECs.
